# When mechanical engineering inspired from physiology improves postural-related somatosensory processes

**DOI:** 10.1101/2022.07.20.500443

**Authors:** Chloé Sutter, Marie Fabre, Francesco Massi, Jean Blouin, Laurence Mouchnino

## Abstract

Being the first stimulated by the relative movement of foot skin and the underneath moving support surface, the plantar tactile receptors (i.e., mechanoreceptors) play an important role in the sensorimotor transformation giving rise to a postural reaction. In this light, a biomimetic surface, i.e., complying with the characteristics of the mechanoreceptors and the skin dermatoglyphs (i.e., pattern of the ridges) should facilitate the cortical processes in response to the somatosensory stimulation involved in the balance recovery motor control. Healthy young adults (n = 21) were standing still either on a biomimetic surface or on two control surfaces (i.e., grooved or smooth), when a sudden but low acceleration of the supporting surface along the lateral direction was triggered. A shorter and more robust evoked somatosensory response (i.e., SEP) was observed when participants were standing on the biomimetic surface. As well, a lower oscillatory response in the theta (5-7 Hz) time-frequency domain in the left posterior parietal cortex (PPC) was observed with the biomimetic surface. The greater shear forces induced by the interaction between the feet and the biomimetic surface during the platform motion was likely at the origin of the increased SEP. Besides, the decrease of theta power suggests that the balance task became less challenging. This interpretation was tested in a second experiment by adding a cognitive task, which should be less detrimental for the postural reaction when standing on a biomimetic surface. Consistent with this hypothesis, a more efficient postural reaction (i.e., shorter latency and greater amplitude) was observed when the cognitive task was performed while standing on the biomimetic surface.

## Introduction

During everyday life, unpredictable circumstances can challenge our equilibrium in different directions while standing. This occurs for example when standing passengers are subjected to unexpected acceleration and braking manoeuvres in public transport. Exquisitely compliant, the skin of the foot sole is deformed well before the passenger’s reaction to the driver’s manoeuvres. This skin deformation is due to the mechanical interaction (i.e., shear forces) generated by the surfaces in contact (i.e., the skin of the feet and the supporting surface) and the gravity force acting on the body mass (i.e., body weight). Although the shear forces are small relative to the weight force during natural quiet standing, they are readily detectable by the tactile receptors (Morasso & Schieppati, 1999). These shear forces, and the consequent skin transient deformations, activate the mechanoreceptors located in the skin allowing the brain to identify the direction and amplitude of the perturbation, before being detected by other sensory inputs (e.g. visual, vestibular, or proprioceptive, Mouchnino & Blouin, 2013). These inputs contribute to shape the postural responses during balance perturbations according to the identified limits of postural stability (Carpenter et al., 2010; Latash et al., 2003; Mochizuki et al., 2006; Murnaghan et al., 2011; Riley et al., 1997).

The low perceptual threshold for detecting horizontal acceleration of the support surface (e.g., down to 0.14 m/2 in Mouchnino & Blouin, 2013) suggests a great responsiveness of the tactile sensory system. This responsiveness could have a twofold origin. First, it could stem from the richness of the receptor types (fast and slow adaptive, type I or II for the main tactile receptors) and from the characteristics of the receptors’ receptive fields (round to oval in shape, extended or small, with sharp or blurred boundaries) (Kennedy & Inglis, 2002). It could also stem from the great compliance (i.e., deformation) of the skin in which the receptors are embedded, which depends, among others, on the footprint (epidermal ridges) orientation. The different fingerprint orientations, relative to the textured surface, enhance the subsurface strain and transmission of tactile information for any direction of the shear forces (Fearing & Hollerbach, 1985; Prevost et al., 2009). The mechanoreceptors and footprint spatial characteristics therefore optimise skin deformation, neural encoding of this deformation and transmission of cutaneous sensory inputs.

While the interaction between a surface and the skin has been extensively studied during finger exploration of a surface (e.g., Hollins & Risner, 2000; Lederman & Klatzky, 2009; Camillieri et al., 2018; Massimiani et al., 2020; see Basdogan et al., 2020 for a review), intriguingly, most of the investigations in the field of balance control have ignored the surface/body contact mechanics. This is particularly surprising given that the foot soles show footprints (dermatoglyphs) that have similar types and density of forms as fingerprints (e.g., ∼60% of loops are shared by fingerprint and footprints, Sarma, 2020). Skin deformation during tactile exploration depends not only on the morphological, topographical, and mechanical properties of the skin, but also from the physio-chemical properties of the surface (i.e., materials, topographic features such as height differences, adhesion, spatial period, Cornuault et al., 2015). In this light, one can hypothesise that the processing of foot cutaneous inputs could be enhanced when standing on a biomimetic surface, whose texture is inspired by the spatial characteristics of both mechanoreceptors and dermatoglyphs.

Here, to specifically test this hypothesis, we recorded and compared the amplitude of the cortical evoked somatosensory potential (i.e., P_1_N_1_ SEP) when the participants were standing either on biomimetic or control surfaces, which translated in the lateral direction. Because it witnesses the amount of sensory input processing at the cortical level (Desmedt & Robertson, 1977; Hämäläinen et al., 1990; Mauguière et al., 1997; Salinas et al., 2000; Lin et al., 2003; Case et al., 2016), the amplitude of the P_1_N_1_ SEP component was expected to be greater when the participants stood on a biomimetic surface than on other types of surface (e.g., smooth) *(Experiment 1*).

Moreover, since an efficient sensory processing allows a better detection of balance threats, the use of a biomimetic surface should decrease the cognitive demand associated to equilibrium maintenance during the translation of the support surface. To test this hypothesis, we compared the changes of sensorimotor theta band power (4-7 Hz) evoked by the translation of the biomimetic and control surfaces. Indeed, recent studies have shown that an increased theta power over sensorimotor areas is an electrophysiological biomarker of the increased difficulty of the balance task. For instance, Sipp et al. (2013) found significant increases of theta power in the left sensorimotor cortex before imminent rightward or leftward loss of balance. This localized change of theta power spread afterward over other cortical areas (e.g., anterior parietal and anterior cingulate areas). Similar increase of theta band activity was observed during the preparation of a challenging balance recovery task that required participants to keep the feet-in place and to refrain stepping responses (Solis-Escalante et al., 2019). Increased theta band activity is also observed in the PPC during the transition from a stable to an unstable surface (Hülsdünker et al., 2015; Mierau et al., 2017). This is in line with de Lafuente & Romo (2006) who showed (in Monkey) responses of the superior PPC to tactile stimulation which occurred ∼60 ms following SI responses. Since the PPC is involved in sensory information integration and generates decision-related activity (see Romo et de Lafuente 2012 for a review), this associative cortical area is likely involved in motor recovery response to balance perturbation.

Based on the premise that balance control requires a minimum state of attention and of cognitive resources (Lajoie et al., 1993; Jacobs et al., 2008; Maki & McIlroy, 2007; see Boisgontier et al., 2011 for a review), facilitating the detection of balance instability when standing on a biomimetic moving surface should decrease the attentional demand required for standing steadily. By using a dual task (DT) paradigm (*Experiment 2*) in which participants were involved in a cognitive task while their supporting surface translated sideways, we expected smaller interference between the postural and cognitive tasks when participants stood on a biomimetic surface compared to other surfaces (e.g., smooth). This should result in a better performance in the cognitive task or a sharper postural reaction to the surface translation (i.e., large, short-duration postural reactions, Redfern et al., 2002).

## Methods

### EXPERIMENT 1

#### Participants and task

Fifteen participants (9 women) without any known neurological and motor disorders participated in the experiment (mean age 26 ± 3 years, mean weight 64 ± 10kg). All participants, except two, characterized themselves as right footed. All participants gave their written informed consent to take part in this study, which conformed to the ethical standards set out in the Declaration of Helsinki and which was approved by the CERSTAPS ethic committee.

Participants were requested to stand barefoot with their feet at a natural distance apart on different types of surfaces (see below), fixed in the middle of a movable force platform. They wore a safety harness attached to the structure top. We ensured that the feet position remained the same throughout the experimental conditions. As the morphology of foot (i.e., flat, hollow, standard) can have an impact on body stability (Klein et al., 2008), we verified that none of them had any foot morphological particularities. This was done by measuring the width of the forefoot (i.e., metatarsal band from the first to the 5^th^ toe) and the isthmus width localized in the middle of the foot and connecting the forefoot with the rearfoot. Computing the percentage ratio ((isthmus width)/ (forefoot width) x 100) allowed us to identified hollow feet (<33%), standard feet (33% to 50%) and flat feet (>50%) (Klein et al., 2008). All participants showed standard feet, therefore none of them have been excluded from the analyses.

We used a set-up employed in previous studies for stimulating foot tactile afferents (e.g., Saradjian et al. 2019). A movable force platform is placed on two parallel rails and is held stationary by an electromagnet (Fig. 1A). A cable is attached to the platform and run laterally through a pulley system with a load fixed to its extremity. The platform translation is triggered by deactivating the electromagnet. The load is adapted to the weight of the participants, such that switching off the electromagnet translated the platform very slowly to the right of the participants, without endangering their balance. A triaxial accelerometer (MEMS, model 4630, Measurement Specialities, USA; 1000 Hz) was used to measure the platform acceleration (mean peak acceleration across participants of 41 ± 4 cm.s^-2^).

**Figure 1.**
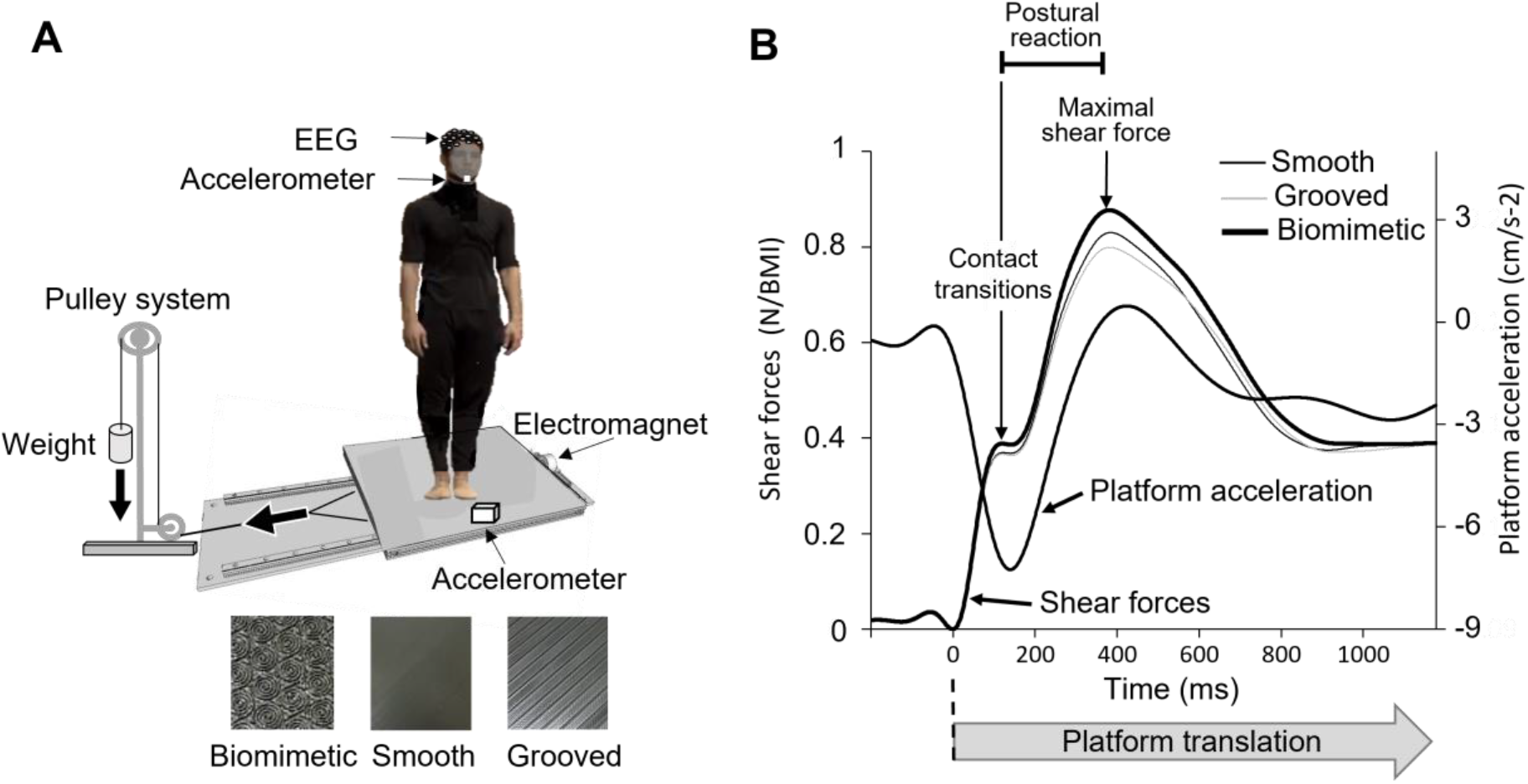
**A**. Experimental set up. The participant stood on one of the three surfaces glued on the force platform which, on deactivation of the electromagnet, would undergo a translation to gravity loading. They wore a safety harness (not shown in the figure) attached to the structure top. **B**. Mean lateral forces and platform acceleration of the 15 participants. At the platform translation onset (broken line) two consecutive phases in lateral force were observed. The first peak force corresponds to the maximal extensibility of the skin under the feet until the frictional force (i.e., shear force) cannot anymore resist the sliding leading to transient variations of the local strain distribution (“skin-surface contact transitions”). Afterwards, a second force peak occurred and corresponds to a postural reaction.

At the start of a trial, the participants looked at fixation point positioned at eye level, 2m directly ahead. They were asked to close their eyes upon receiving the verbal information on the nature of the upcoming condition, and to remain still. This information indicated one of these two conditions: platform translation (37 trials) or platform steady (8 trials), which were pseudo-randomly distributed. The later set of trials reduces the possibility of adopting a stereotyped postural set linked to the forthcoming body translations. These trials were also used to measure and model the noise contaminating the EEG data (see below). In all trials, the participants had to maintain an upright steady posture during 5 s (i.e., duration of trial recording). The platform translation occurred at any time between 2 s and 4 s after providing the information about the platform translation to avoid anticipating the instant of the translation onset. The trials without translation also lasted 5 s. A short break was frequently proposed to the participants during the experiment.

#### Surfaces

Participants were standing on three different surfaces that were glued on the platform: a biomimetic surface, a textured surface with no bioinspired features characteristics (i.e., grooved) and a smooth surface (the last two surfaces were used as controls). These surfaces were created with a 3D printer (Ultimaker 2+) using biopolymer thermoplastic (Polylactic acid, PLA). Three characteristics were selected to build the biomimetic surface: shape, spatial period, and depth of the ridges.

The biomimetic surface was textured with circular or oval shapes inspired from both the shape of the tactile receptors’ fields that demonstrate a preferential skin strain axis and orientation of this axis, which is not the same for all units (Kennedy & Inglis, 2002; Valbo et al., 1995), and the forms of the dermatoglyphs, which exhibit three main circular forms (loops, whorls, and arcs, first described by Cummins & Midlo, 1926). We verified whether the radius of curvature of the circular shape of the biomimetic surface complied with the participants’ toe prints. To do so, we used the ink dabbing method to collect the toe prints of each participant on a white sheet of paper. Contrary to fingerprints, toe prints have their characteristic features in the lower end of the phalanges (Sarma, 2020). Then, the rolling of the prints was taken longitudinally from lower end to the upper end of the toe (i.e., opposite direction than when collecting fingerprints). For each participant, we measured, and then averaged, the radius of curvature of the 3 most visible ridges from 3 different toes. A t-test of means against a reference value indicated that the radius of curvature on the toes surface (4.3 ± 1.1 mm) did not differ significantly compared to the radius of the loops printed of the biomimetic surface (t_13_= 1, p= 0.34).

The spatial period of the biomimetic surface complied with the period of the participants’ toeprints ridges. This was confirmed by the result of the t-test of means against a reference value showing that the mean period of the biomimetic surface (0.9 mm) was not significantly different from the period of the toeprints ridges (0.87 ± 0.06 mm) (t_13_= -1.66, p= 0.12). Note that the spatial period of the biomimetic surface also complied with the distance between the centre of adjacent receptive fields of the mechanoreceptors (from 0.9 to 3.8 mm; Johansson & Vallbo, 1983).

Finally, the depth of the valley was computed from what we know from finger surface exploration and balance maintenance literature. The depth to properly perceive the stimulus on the finger skin is estimated as 0.1 mm with a 0.5 N normal force (Camillieri et al., 2018; Peyre et al., 2017). In a previous study, we found that the minimum shear forces amplitude to detect support translation beneath the feet standing in a natural position was ∼3.5 N (Mouchnino & Blouin, 2013). This lead to the suggestion that a 0.7 mm depth of the valley should enable to perceive the minimal shear force when bearing our body weight.

A smooth surface also printed in PLA but without any designed patterns was used as a control surface. While the smooth surface is used as standard control surface, a textured surface (i.e., grooved surface) with different texture parameters, different from the ones of the dermatoglyphs and characterized by a main direction, allows for excluding a simple effect of the local strain variation due to a general texture. Comparing the biomimetic texture with a standard texture can then highlight the role played by a geometrical distribution of the texture mimicking the receptive features of the foot skin. Such a biomimetic geometry can give rise to local stress and strain distributions with a specific orientation pattern, which can favour the detection of the transient strain variations by the mechanoreceptor activation. The texture of the grooved surface was composed of rectilinear ridges with a depth of 0.3 mm and a period of 7 mm.

The order of the 3 surface expositions was counterbalanced across participants. The participants were not informed prior to the experiment about the reason the standing surface was changed during the recording session. When the participants were asked after the experiment whether they had perceived that they stood on surfaces having different textures, none of them reported having done so.

#### Recordings and analyses

Electroencephalography (EEG) activity was continuously recorded from 64 Ag/AgCl surface electrodes embedded in an elastic cap (BioSemi ActiveTwo system: BioSemi, Netherlands). Specific to the BioSemi system, “ground” electrodes were replaced by Common Mode Sense active and Driven Right Leg passive electrodes. The signals were pre-amplified at the electrode sites, post amplified with DC amplifiers, and digitized at a sampling rate of 1024 Hz (Actiview acquisition program). The signals of each electrode were referenced to the mean signal of all electrodes. Four Ag/AgCl electrodes placed near the canthus of each eye and under/over the left eye orbital allowed us to control for blinks, and horizontal and vertical eye movements.

The continuous EEG signal was segmented into epochs synchronized relative to the onset of the platform translation, which was identified at the onset of the monotonic increase of the shear force. After artefact rejections based on visual inspection, for each participant and surface, a minimum of 96% of the trials were included in the analyses. The signals were filtered off-line with a 50 Hz digital notch filter (24 dB/octave) and with a 0.1–48 Hz band-pass digital filter (48 dB/octave) implemented in BrainVision Analyzer 2 software (Brain Products, Germany). For each participant, the SEPs were obtained by averaging all epochs of each surface for each participant. The average amplitude computed 50 ms prior to the platform translation served as baseline. Consistent with studies recording cortical potentials evoked by lower limb stimulation (Altenmüller et al., 1995, Saradjian et al., 2013), the SEPs were found to be maximal over the vertex (Cz electrode). Therefore, this electrode was used to assess the SEPs. We primarily based our analyses on the P_1_N_1_ wave evoked by the sensory stimulation induced by the platform translation. The amplitude of P_1_N_1_ was measured peak to peak, and its latency was assessed measuring the P_1_ latency.

#### Cortical sources

Neural sources of the SEPs were estimated with the dynamical Statistical Parametric Mapping (dSPM, Dale et al., 2000) implemented in the Brainstorm software. A boundary element model (BEM) with three realistic layers (scalp, inner skull, and outer skull) was used to compute the forward model on the anatomical MRI brain template from the Montreal Neurological Institute (MNI Colin27). Using a realistic model has been shown to provide more accurate solution than a simple three concentric spheres model (Sohrabpour et al, 2015). We used of a high number of vertices (i.e., 15 002 vertices) to enhance the spatial resolution of the brain template. Such EEG source reconstruction has proved to be suited for investigating the activity of outer and inner cortical surfaces with 64 sensors (Chand & Dhamala, 2017; Ponz et al., 2014). Measuring and modelling the noise contaminating the EEG data is beneficial to source estimation. Noise covariance matrices were computed using the 8 trials with the platform steady condition, while the participants stood still. The current maps were averaged from the start of the shear forces to N1, for each participant and surface condition.

The data were transformed into time-frequency domain using Morlet wavelet transforms. We used a 1 Hz central frequency (full width at half maximum FWHM tc=3sec) which offers a good compromise between temporal and spectral resolutions (Allen & MacKinnon, 2010). The power of theta (5-7 Hz) was computed for each trial in the source space in a region of interest (ROI, 589 vertices) encompassing the left inferior and superior PPC (based on the Destrieux cortical atlas, Destrieux et al., 2010). Then, the signal was expressed as a change of theta power computed over the first 400 ms of the platform translation (which included the P1N1 SEP) with respect to a 350 ms window baseline taken before the translation (−400 to -50 ms). For each participant, the resulting event-related synchronization/desynchronization (ERS/ERD) was then averaged across trials and surface conditions. The same procedure was applied with the signals computed in a control ROI (650 vertices) encompassing the inferior and superior PPC of the right hemisphere.

#### Behavioural recordings and analyses

The ground reaction forces and moments were recorded with an AMTI force platform (60 × 120 cm, Advanced Mechanical Technology Inc., USA) at a sampling rate of 1000 Hz. The ground reaction shear forces were analysed along the mediolateral (M/L) axis, as they represent the earliest signature of the cutaneous stimulation evoked by the platform translation. The onset of this stimulation was defined as the first time the M/L shear force started to increase monotonically. Figure 1B shows and the dynamics of the M/L shear forces resulting from the platform acceleration.

A general pattern of events emerged:

i. the first phase shows a ramp of the platform acceleration (constant jerk) corresponding to a ramp in the traction force; this phase lasted on average 197 ± 11ms leading to a platform displacement of about 2.93 ± 0.30 mm. Due to the inertia of the body, this displacement is accommodated mainly by the skin deformation. The peak amplitude of this force was measured with respect to its baseline value (i.e., prior to the translation onset);
ii. in a following transition phase, a low valley was observed; the jerk decreases and becomes negative. This is a crucial phase, where the contact between the skin and the surface is characterized by high superficial shearing, leading to transient variations of the local strain distribution (“*skin-surface contact transitions*”), which are directly affected by the topography of both the skin and the surface itself; moreover, local detachments and slipping can occur, leading to transient deformations and waves propagating in the skin, which are likely to activate the mechanoreceptors (“*contact stimuli*”); a similar phenomenon has been observed in literature, when considering the motion of a surface under a stationary fingertip, showing that the deformation of the skin increases until the frictional force (i.e., shear force) cannot anymore resist the sliding (i.e., stick/slip phenomenon, Rabinowicz 1956); within this phase, the shear forces result from a balance between inertia effects and contact accommodation.
iii. Afterwards the shear forces continued to increase in the same direction until a second peak was reached before reversing the forces. This second increase can be considered as a postural reaction (Lhomond et al., 2021; Saradjian et al., 2019). The duration of the postural reaction was defined as the time elapsed between these peaks.

The amplitudes of both peaks of the shear forces were normalized to the body mass index (BMI) of each participant.

Head acceleration was measured with a triaxial accelerometer (model 4630, Measurement Specialities, USA; 1000 Hz) placed on the participants’ chin. We measured head acceleration to evaluate the latency of the vestibular stimulation induced by the platform translation. The onset of the vestibular stimulation was defined at the first moment head acceleration exceeded 0.048 m.s^-2^ (i.e., threshold for vestibular stimulation, Gianna et al. 1996). We measured the lag between the head acceleration and the shear forces to determine if the vestibular stimulation occurred after the stimulation of the foot mechanoreceptors.

Bipolar surface electromyography (EMG, Bortec AMT-8 system, Bortec Bomedical, Canada) was used to record the activity of the long fibular muscle (FL) of both legs. The FL muscles are responsible (with other muscles) for controlling stance. They contribute to the eversion movement of the foot, and also to the maintenance of the arch of the foot to ensure optimal postural stability (Pietrosimone & Gribble, 2012; Jeon et al., 2021). The FL EMG signals were pre-amplified at the skin site (×1000), sampled at 1000 Hz, band pass filtered from 20 to 250 Hz, and rectified. Two participants were excluded from the EMG analyses due to noisy EMG signals. To quantify the muscle activity, we computed the integral of the EMG activity (iEMG) over two intervals. The first corresponded to the “resting interval”. It lasted 1s and ended at the onset of the platform translation. The second interval covered the time elapsed between the platform onset and the N1 component of the SEP. The duration of this second period was specific to each participant. In order to be able to compare muscle activities between the two intervals, we normalized, for each participant, the EMG activity of the resting interval to the duration of the second interval “N1 latency” (∼180ms). We also calculated the latencies of EMG changes relative to the onset of the platform translation. This was done by first computing the mean and standard deviation of the muscle background activity (i.e., during the resting interval) for each participant and surface. The onsets of the EMG increased, or decreased activities were defined as the times at which the EMG activity increased above or decreased below a threshold level set at twice the standard deviation of the mean background activity.

### EXPERIMENT 2

#### Participants and task

The goal of *Experiment 2* was to test whether the attentional demand required for equilibrium maintenance is reduced when standing on the biomimetic surface, compared to a control surface. To further compare the effect between standing on a biomimetic and standing on a non-biomimetic textured surfaces, the grooved surface used in *Experiment 1* was selected as the control surface.

Twenty-one new participants (7 women) without any known neurological and motor disorders participated in the experiment (mean age 22 ± 2 years, mean weight 67 ± 10kg). All participants gave their written informed consent to take part in this study, which conformed to the ethical standards set out in the Declaration of Helsinki and which was approved by the CERSTAPS ethic committee.

The procedure was identical to that in *Experiment 1* with one exception that pertained to the dual task (DT) paradigm. We used a demanding cognitive task to increase the participants’ attentional load while their supporting surface was translated as in *Experiment 1*. Participants were asked to listen to a series of 10 different three-digit numbers (1Hz) that ended 60ms before the platform translation. The 10 numbers were spelled out at high speed (being completed in 10 s) by a computer voice. The series of numbers varied across trials but were the same for both surface conditions and participants. Participants were instructed to count silently the number of times that 7 was part of the three-digit numbers and to provide their response at an auditory tone occurring 3 s after the platform translation onset (i.e., after the data analysed intervals). The same procedure was used for the trials in the steady platform condition. We counterbalanced the presentation of the different conditions (i.e., biomimetic or grooved surfaces, with or without translation; single or dual tasks) across participants but prevented the occurrence of 2 successive conditions involving the dual task.

The participants’ performance in the cognitive task was assessed by computing, for each surface condition, the averaged percentage of errors ((number of errors/10 numbers) x 100). An error was counted each time that the participant reported a wrong number of times that 7 was part of the spelled-out numbers.

#### Statistical analyses

The behavioural and EEG data were submitted to separate analysis of variance (ANOVA) with repeated measurements. In *Experiment 1*, one-way ANOVAs were used for mean comparisons with the support surface (Smooth, Grooved, Biomimetic) as intra-participants factor. We computed statistical maps by contrasting the current maps (i.e., each vertice) computed when standing on a biomimetic surface and control surfaces using t-tests (significance threshold p < 0.05) (Tadel et al., 2011). We applied an FDR (False Discovery Rate) correction for multiple comparisons across regions (Benjamini & Hochberg, 1995).

In *Experiment 2*, a 2×2 ANOVA was used for mean comparisons with the support surface (Grooved, Biomimetic) and task (single or dual task) as intra-participants factors. Significant effects (statistical threshold of p ≤ 0.05) were further analysed using Newman-Keuls post-hoc tests.

### Experiment 1

#### Results

##### Augmented peripheral stimulation by the biomimetic design of the surface

As shown in Fig. 1B, the shear forces increased in the leftward direction at the onset of the rightward platform translation until a clear break down point was reached (see methods). The results showed that the amplitude of this early force differed significantly between surfaces (Fig. 2B; F_2,28_ = 13.84, p < 0.05). Post-hoc analyses revealed that the amplitude was greater for the biomimetic (0.40 ± 0.05 N/BMI) than for the grooved and smooth surfaces (Ps < 0.05), which did not differ significantly (p = 0.20; overall mean of 0.38 ± 0.05 N/BMI).

**Figure 2.**
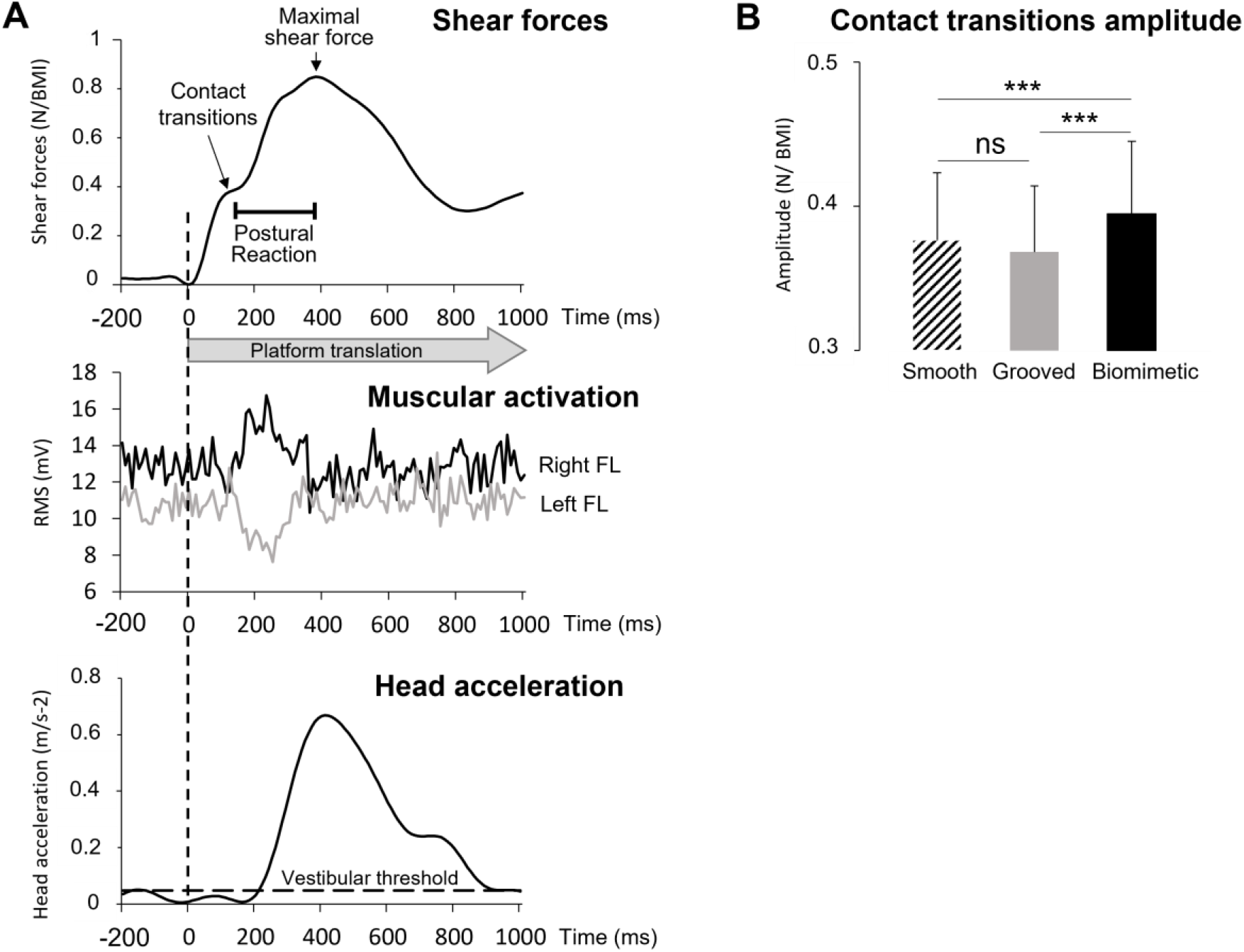
**A**. Averaged shear forces (top panel), muscular activities (middle panel) and head accelerations (bottom panel) for all trials of one typical participant standing on the grooved surface. The time 0 (broken line) corresponds to the onset of the platform translation. **B**. Mean contact transitions amplitudes for all 15 participants computed on the three surfaces (smooth, grooved, biomimetic). Error bars represent standard deviation across participants; *** p < 0.001, ns = not significant.

The latency of the break down point did not depend on the surface (overall mean of 117 ± 12ms; F_2,28_ = 0.12, p= 0.89). It occurred before reaching the maximal value of the platform acceleration (overall mean -26 ± 11ms) for all surfaces (no significant surface effect F_2,28_ = 2.09, p= 0.14); this suggested that the early peak of the shear forces was not generated by the reaching of the maximum acceleration of the platform and the consequent change in the sign of the jerk.

In addition, the EMG analyses (Fig. 2A) showed that the activity of the right FL muscle started to increase 40 ± 20 ms after the break down point and that the latencies of these increases were not significantly affected by the surface (F_2,24_= 0.64; p = 0.54). The left FL activity decreased simultaneously with the right FL activation in 10 out to 13 participants. The delay of the changes in the FL muscles activities relative to the break down point suggested that the early shear forces were not muscularly induced, but rather passively/mechanically evoked by the contact transmission between the stretched skin and the platform.

Besides, the observations that only the right FL (among the 2 recorded muscles) started to be activated after the “skin-surface contact transitions” (i.e., during the postural reaction, Fig. 2A) suggested that it was engaged in the breaking of balance perturbation. The postural reaction was not altered by the surface conditions, neither for its amplitude (F_2,28_= 2.50; p = 0.10, mean = 0.53 ± 0.10 N/BMI) nor for its duration (F_2,28_= 0.20; p= 0.82, mean = 296 ± 64ms). The head started moving (i.e., accelerate) during the postural reaction with a latency of 172 ± 38 ms relative to the platform translation (Fig. 2A). This lag, which likely resulted from the body mass inertia, was not significantly affected by the surface condition (F_2.28_ = 1.48; p= 0.25).

##### Cortical facilitation of sensory input when standing on a biomimetic surface

To determine whether the SEP (i.e., P_1_N_1_) originated from tactile and/or vestibular peripheral inputs, we compared the latencies of P_1_ and N_1_ to the vestibular stimulation onset. The onset of the vestibular stimulation was defined at the first moment head acceleration exceeded 0.048 m.s^-2^ (i.e., threshold for vestibular stimulation, Gianna et al. 1996). This threshold latency was not significantly affected by the surface condition (F_2,28_= 1.75; p = 0.19).Paired t-tests showed that P_1_ and N_1_ latencies significantly preceded vestibular stimulation onset for all the surfaces (see Table 1). This indicated that the SEP was not evoked by vestibular inputs, but more likely by cutaneous inputs originated from the shear forces (i.e., skin strain) evoked by the platform translation.

**Table 1.** Mean latencies of all participants (n = 15) and inter participants standard deviation (SD) for the P1, N1 and the time when the head reached the vestibular threshold as a function of the surfaces on which participants were standing. The paired t test corresponds to the comparison between the P1 or N1 and the vestibular threshold.

| Smooth |  | Grooved |  | Biomimetic |  |
| --- | --- | --- | --- | --- | --- |
| Vestibular threshold |  |  |  |  |  |
| 211ms (±40) |  | 225ms (±27) |  | 207ms (±32) |  |
| P1 | N1 | P1 | N1 | P1 | N1 |
| 138ms (±11) | 186ms (±15) | 137ms (±13) | 184ms (±18) | 126ms (±16) | 187ms (±13) |
| T test |  |  |  |  |  |
| t = 7.09 ; p<0.001 | t = 2.79 ; p = 0.01 | t = 12.12; p<0.001 | t = 5.50; p<0.001 | t = 9.93; p<0.001 | t = 2.68 ; p = 0.02 |

Importantly, the latency of P_1_ was shorter and the amplitude of the P_1_N_1_ was greater for the biomimetic surface than for the smooth and grooved surfaces (Fig. 3). The significant main effect of the surface condition (contact stress and strain distribution obtained by the biomimetic topography) was confirmed by the P_1_ latency (F_2.28_ = 6.65, p = 0.004) and the P_1_N_1_ amplitude (F_2,28_ = 3.55, p = 0.04), which did not show difference between the two control surfaces either for P_1_ latency (p= 0.81) or P_1_N_1_ amplitude (p= 0.65). There was no significant effect of surface on N1 latency (F_2.28_= 0.41, p= 0.67).

**Figure 3.**
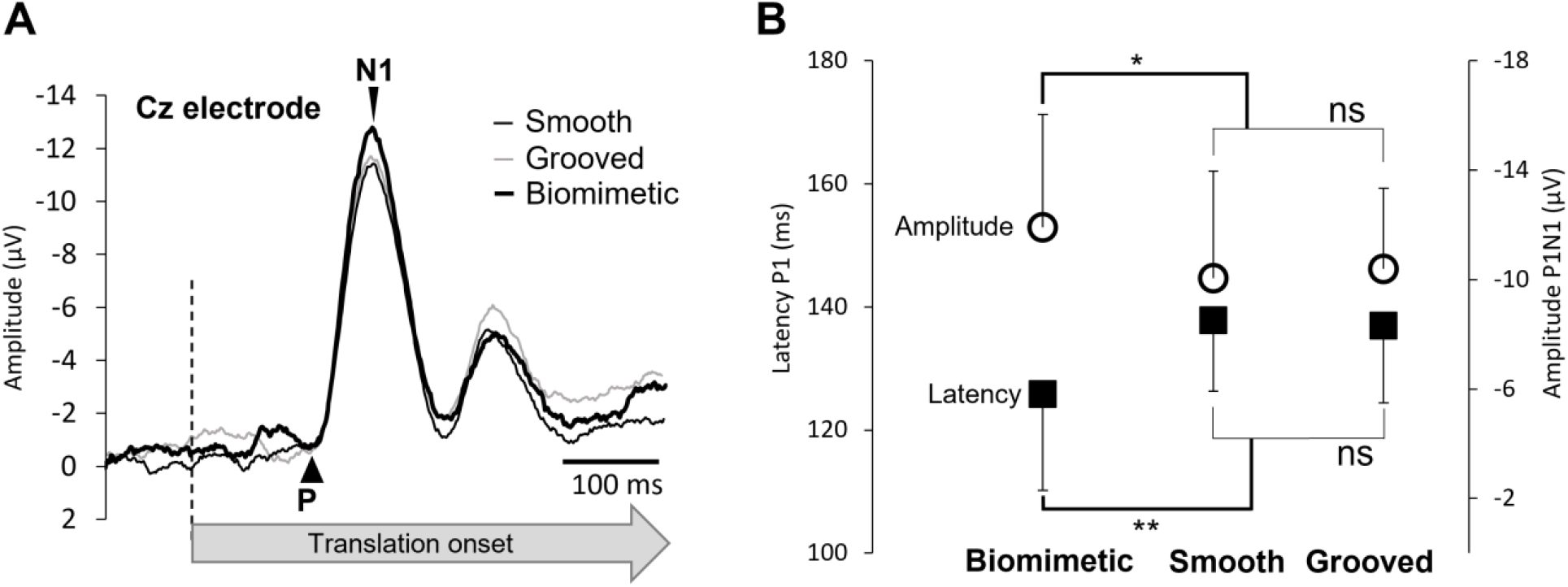
**A**. Grand average (n=15) of the SEP recorded over Cz electrode for the 3 surfaces (biomimetic, smooth, grooved). The broken line indicates the moment of the stimulation (i.e., translation onset). **B**. Mean P1 latency and amplitude of the averaged P1N1 SEP for all participants on the three surfaces (biomimetic, smooth, grooved). Error bars represent standard deviation across participants, *p < 0.05; **p<0.01; ns = not significant.

To verify if the SEP facilitation (i.e., shorter P_1_ latency and greater P_1_N_1_ amplitude) observed with the biomimetic surface could be linked to a change in leg muscle activity, we compared the iEMG of the right and left FL during the N_1_ latency interval. The ANOVA did not show a surface condition effect on the iEMG (F_2.24_= 0.57; p = 0.57 and F_2.24_= 1.11; p = 0.34, for the right and left FL, respectively). These results suggest that the changes in the SEP observed over the somatosensory cortex when the participants stood on the biomimetic surface stemmed from an increased afferent volley from the foot sole mechanoreceptors rather than from an altered motor command (i.e., muscle activity).

##### Standing surface-specific source localization

The statistical cortical maps revealed significant greater activation (i.e., current) in the left precuneus (medial extent of Brodmann area 7, warm colour) for the biomimetic surface for both the biomimetic/smooth (Fig. 4A) and biomimetic/grooved (Fig. 4B) contrasts. The same contrasts also showed significantly greater activity of the extrastriate body areas (EBA, BA19). On the other hand, these contrasts revealed greater activities of the left premotor (PM) and anterior cingular (ACC) cortices (see cold colors in Fig. 4A) for the smooth surface and greater activation of the right inferior PPC (BA39) for the grooved surface (Fig. 4B, cold colour).

**Figure 4.**
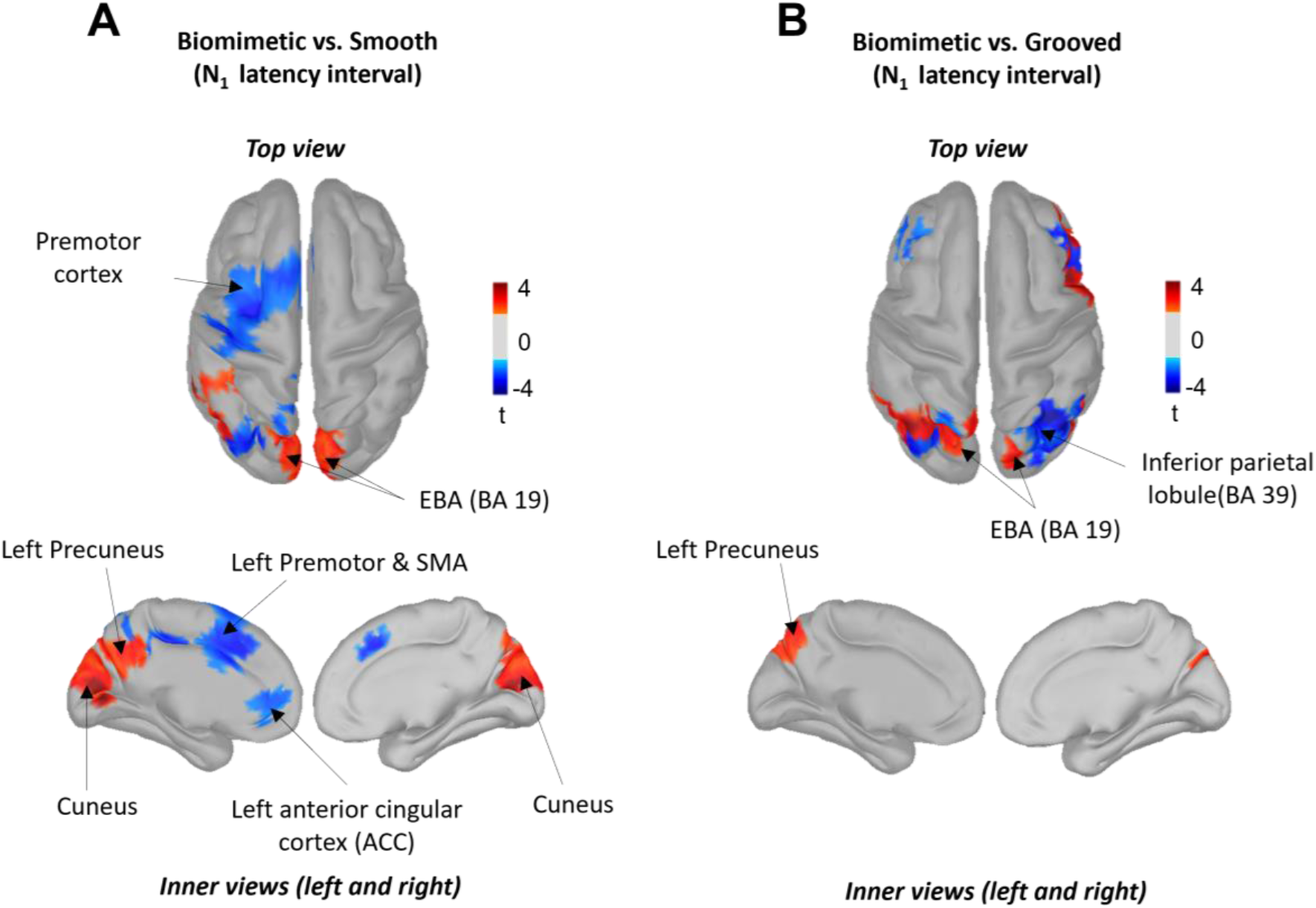
Statistical source estimation maps for Biomimetic versus Smooth (A), Biomimetic versus Grooved (B) contrasts. Significant t-values (p ≤ 0.05, n = 15) of the source localization were shown during the time window from 0 ms to N1 latency. Sources are projected on a cortical template (MNI’s Colin 27). For each contrast, we display the top and the left and right inner cortical views.

##### Modulation of theta (5-7 Hz) oscillations in the left PPC

The time-frequency analysis showed that theta power computed in the left PPC was significantly modulated by the type of surface on which the participants were standing (F_2.28_ = 3.99; p = 0.03). Post-hoc analyses revealed that theta power was significantly smaller for the biomimetic than for the smooth and grooved surfaces (ps < 0.05), with no significant difference between the two control surfaces (p= 0.67, ps > 0.05) (Fig. 5C). This effect was lateralized to the left hemisphere as it was not observed in the right PPC (F_2.28_ = 1.32; p = 0.28).

**Figure 5.**
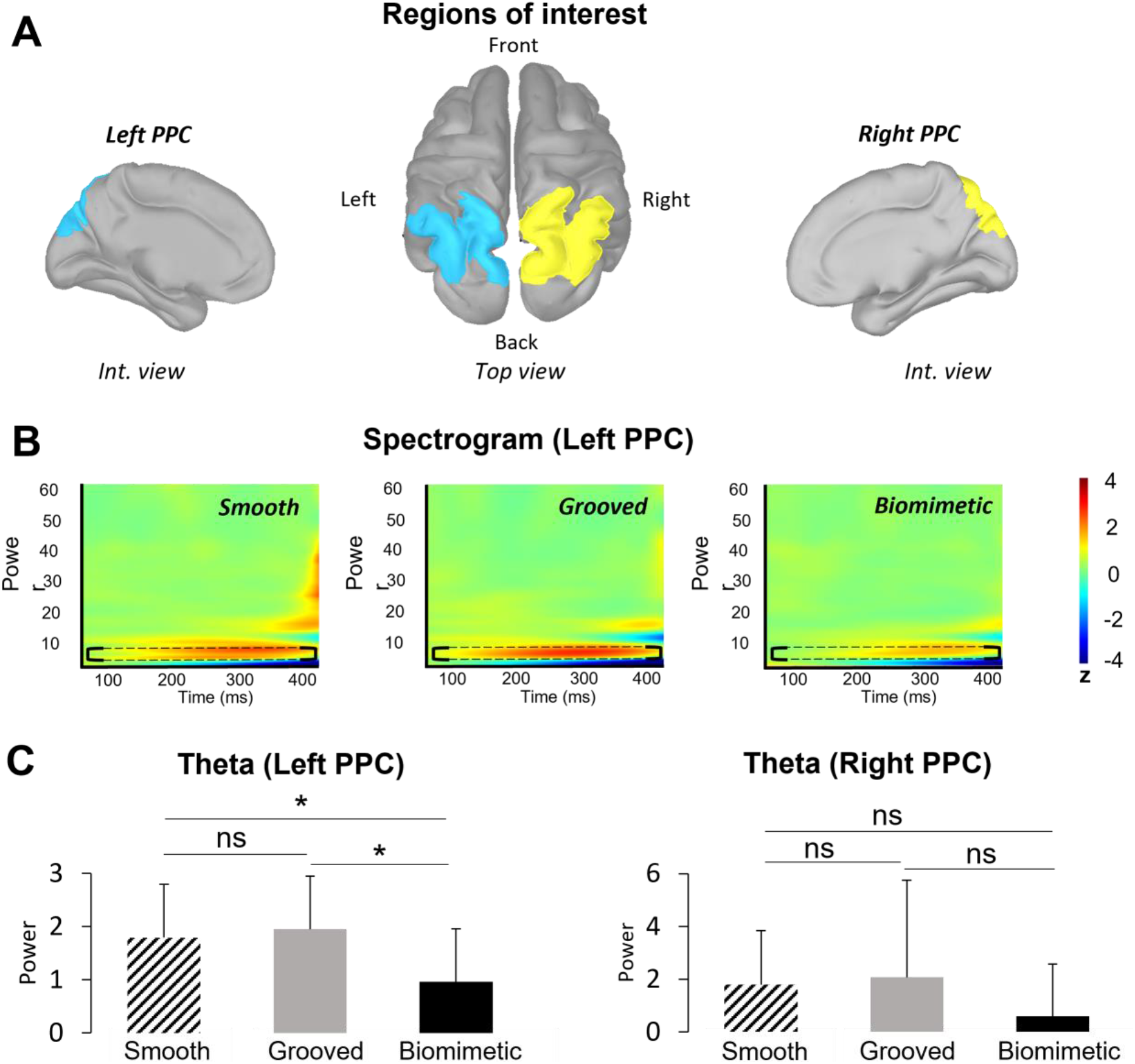
**A**. Localization of the regions of interest (ROIs) on the anatomical MRI Colin 27 brain template that was used to compute cortical activations. Note that similar ROIs were defined for the left and right Parietal Posterior Cortex (PPC). **B**. Time-frequency power (ERS/ERD) of the signals by means of a complex Morlet’s wavelet transform applied on the ROIs for each surface of each participant then averaged. Cooler colors indicate ERD and warmer colors, indicates ERS. Frequency bands from 1 to 90 Hz were provided to have an overview of the full spectral content of cortical neural oscillations. We showed the spectrum from 0 to 400 ms to focus on the analyzed time window of the ERS/ERD (thereby removing edge effects). The theta band has been circled in dotted line for each surface. **C**. Mean of theta (5-7 Hz) frequency band computed during (0; 400 ms) time window in the left and right PPC for each surfaces (smooth, grooved, biomimetic). Error bars are standard error across participants, * p < 0.05, ns = not significant

## Discussion

### Standing on a biomimetic surface speeds up and enhances the sensory transmission from foot to cortical areas

Complying with both dermatoglyphs and mechanoreceptors characteristics, the biomimetic surface in interaction with the feet evoked faster and greater cortical response to the platform translation (i.e., P_1_N_1_), with respect to both control surfaces (i.e., grooved and smooth). The greater amplitude of the SEP, when standing on the biomimetic surface, suggests augmented cutaneous afferent processes (see Desmedt & Robertson, 1977; Hämäläinen et al., 1990; Salinas et al., 2000; Lin et al., 2003; Case et al., 2016; Lhomond et al., 2016). The increase P_1_N_1_ amplitude and the shortening of the evoked response latency (i.e., P_1_) could be due to an accentuation of the deformation of the skin when interacting with the biomimetic surface. The enhanced skin transient deformations was witnessed by an increase amplitude of the peak force during the “skin-surface contact transitions” while no concomitant change of its duration was observed. This is in line with Tang et al.’s (2020) study showing that a greater deformation of the fingers’ skin generates greater friction force, and stress that induce stronger tactile stimulation of the mechanoreceptors. Similarly, the interaction of the foot and the biomimetic surface increased the intensity of the skin mechanoreceptors stimulation, which in turn boosted the transmission of cutaneous signals to the primary somatosensory cortex (SI) where neurons respond to various tactile stimulations (e.g., Bensmaia et al., 2008; Weber et al., 2013).

Our source analyses showed that changing the standing surface significantly altered the current recorded in the left PPC but had no significant effect on the current recorded in SI. This suggests that the sensory facilitation observed with the biomimetic surface may have involved direct thalamocortical projections to the PPC. The basis for this suggestion is twofold. First, neuroanatomical studies in the macaque have shown direct projections of cutaneous information from the thalamus to the PPC (Pearson et al., 1978; Padberg et al., 2009; Impieri et al., 2018; Gamberini et al., 2020). Secondly, our results showed an increased activation of the medial extent of the SPL (i.e., precuneus) for the biomimetic surface, with respect to both control surfaces, in line with functional interactions (e.g., somatomotor function) between the precuneus and the thalamus (Cunningham et al., 2017; Gamberini et al., 2020). In this light, the shared connectivity between the precuneus and the extrastriate body area (EBA) (Zeharia et al., 2019), which showed an enhanced activity in the biomimetic surface condition, is consistent with the role of the EBA in enhancing the local spatial processing of body information on the direction of a stimuli out of view (Striem-Amit & Amedi, 2014; Urgesi et al., 2007), such as the skin stretch under the feet in the current study.

The shorter latencies of the SEP observed when the participants stood on the translating biomimetic surface could suggest some contribution of muscle-stretch receptors, as it is well-established that muscle spindle endings are extremely sensitive stretch receptors (see Starr et al., 1981; Cohen et al., 1985). However, the evoked potentials that arise from the stimulation of the lower limbs endings exhibit much shorter latencies (i.e., 20-65 ms; Starr et al., 1981; Cohen et al., 1985) than those observed in the current study (126 ms, for the biomimetic surface). Furthermore, the increased activity of the leg muscles induced by the translation of the biomimetic surface occurred ∼160 ms after the onset of the translation, i.e., after the P1 occurrence. This reduces the possibility for a significant role of muscle-stretch receptors in generating the SEP when the participants stood on the translating surface, irrespectively of the type of supporting surface. Neither is vestibular input a likely candidate for evoking the cortical response, as P1 and N1 had shorter latencies than the latency with which the head reached the acceleration threshold for activating vestibular receptors (i.e., 207 ms, for the biomimetic surface).

### Decreased balance task difficulty when standing on the biomimetic surface

Finally, the results of theta oscillations analyses were consistent with the likely role of the left PPC in attentional processes (Hülsdünker et al., 2015; Mierau et al., 2017). Our participants showed a significant decreased power of theta oscillations in the left PPC when they stood on the biomimetic surface compared to the control surfaces (i.e., grooved and smooth). Since theta oscillations are considered as a neural correlate of a need for attentional demand in challenging balance tasks (i.e., theta power increases with increased attentional demand, Sipp et al., 2013; Hülsdünker et al., 2015; Gebel et al., 2020), the significant decrease of theta-band power with the biomimetic surface may reflect a decrease in the attentional demand and a down modulation in the difficulty of the task (Vuillerme & Nafati, 2007). Alternatively, the increased theta power, observed when standing on both control surfaces, may witness the increase attentional and cognitive demands. This is consistent with the greater activities observed within the pre-motor cortex (e.g., SMA) and ACC of the left hemisphere observed when the tactile salience of the surface decreases, as when standing on a smooth surface (as compared to either the biomimetic or grooved surfaces). Previous studies have suggested that the SMA plays an important role in the control of demanding balance tasks (Taubert et al., 2010, 2011). This was notably evidenced by the significant structural and functional adaptation of the SMA activity after balance training (Taubert et al., 2011). Therefore, the enhanced activity of the SMA, found when individuals stood on the smooth surface, may suggest that the standing task was more demanding with this surface than with textured surfaces (biomimetic or grooved). The increased activity observed in the ACC would also be consistent with this suggestion as the role of this cortical region is well-recognized when individuals are uncertain about fulfilling the required task appropriately (e.g., Gemba et al. 1986) or in error-recognition (Holroyd et al., 1998; see Holroyd & Coles, 2002, for a review). These interpretations are also supported by the proposed function of the ACC in the regulation of attention and cognitive control (Botvinick et al., 2001; Bryden et al., 2011; Petersen & Posner, 2012). Enhancing the need for attention when standing on a smooth surface could be a mean for withholding potentially erroneous responses in conditions with impoverished tactile cues on platform translation until other sensory modalities (e.g., vestibular, visual) can resolve the ambiguity of the support displacement. In addition to the increased theta oscillations power for the control grooved surface, the source analyses revealed an increased activation of the right PPC. This findings are consistent with the crucial role of this cortical area in the processing of somaesthetic gravitational information for postural control, as shown in neglect patients after right hemispheric strokes (Pérennou, 2006).

Overall, our results point to a reduced difficulty of the balance task when standing on a biomimetic surface. In *Experiment 2*, we tested the hypothesis that standing on such a surface benefits postural control when the balance task is challenged by increasing the attentional load Indeed, because postural control is known to requires attention (Lajoie et al., 1993), therefore the postural perturbation observed when performing a simultaneous cognitive task would be due to the sharing of limited attentional resources (Stelmach et al., 1990; Chen et al., 1996; Shumway-Cook & Woollacott, 2000; see Woollacott & Shumway-Cook, 2002 for a review).

### Experiment 2

#### Results

##### Cognitive task performance

To verify whether the participants’ performance in the cognitive task differed when standing on the moving biomimetic and the grooved surfaces, we analysed the averaged percentage of errors reporting the number of times that 7 was part of the series of 3-digit numbers spelled out with high speed. A paired t-test did not reveal significant difference in the percentage of errors between the biomimetic and the grooved surfaces (t_20_= -0.20; p = 0.85; 24 ± 11.5%). As shown in Fig. 6A, 2 out 21 participants exhibited values three times above the standard deviation of the mean (i.e., 59% and 66% of errors). These large errors suggest that the task was too difficult for these participants or that they did not allocate enough attentional resources to the cognitive task. These participants were excluded from the analyses.

**Figure 6.**
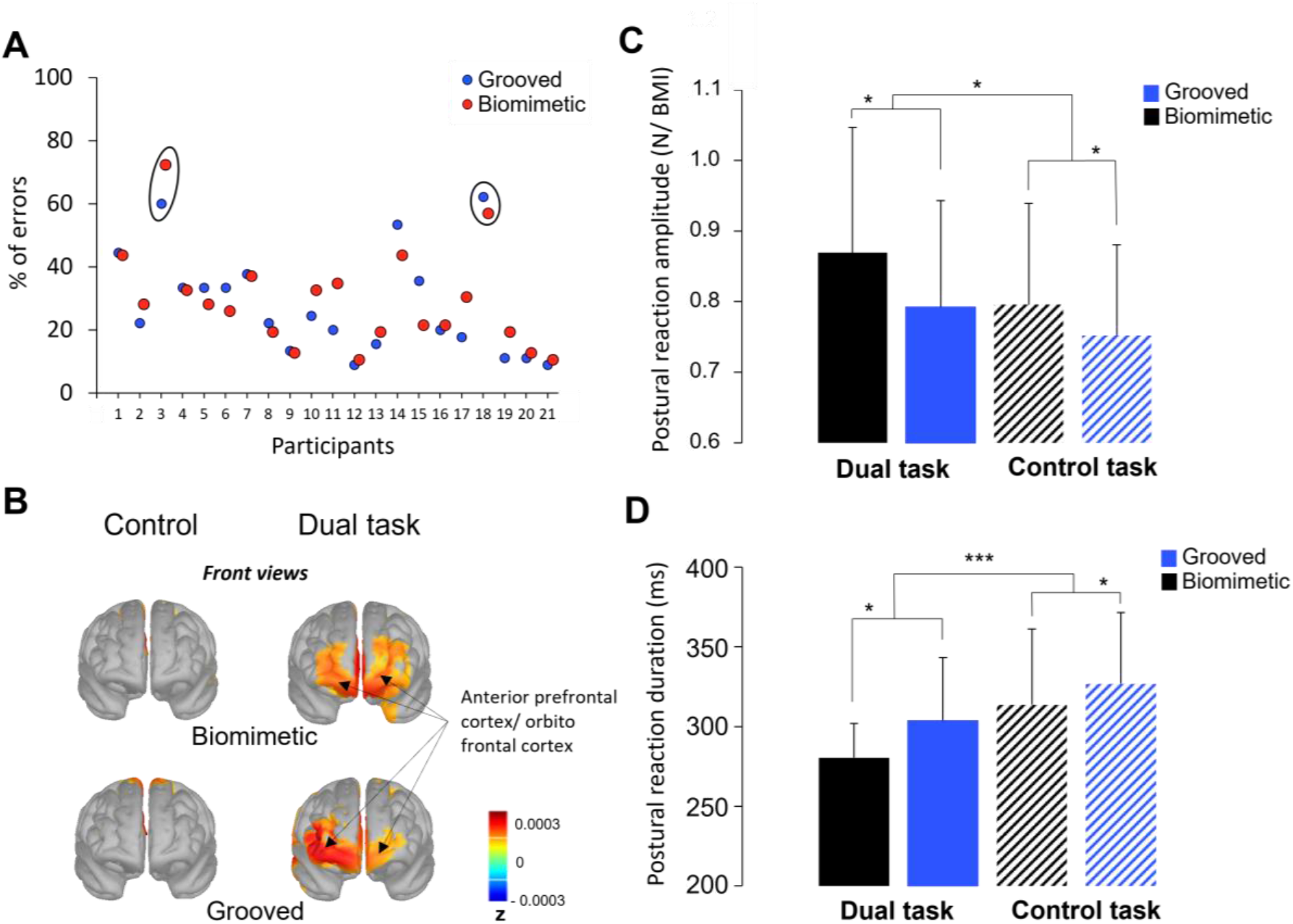
**A**. Percentage of errors for the cognitive task of each participant on both surfaces (grooved, biomimetic). **B**. Source localization during the time window from -2000 to 0 ms latency interval on both surfaces (grooved, biomimetic) during solely the motor task (i.e., control) and the dual task. We display the front view of the sources are projected on a cortical template (MNI’s Colin 27). **C**. Mean postural reaction amplitude normalized by the BMI for all participants computed on both surfaces (grooved, biomimetic) during solely the motor task (i.e., control) and the dual task. Error bars represent standard deviation across participants; * p <0.05) **D**. Mean postural reaction latency for all participants (n = 21) computed on both surfaces (grooved, biomimetic) during solely the motor task (i.e., control) and the dual task. Error bars represent standard deviation across participants; *** p < 0.001).

Furthermore, the activity observed over both the anterior prefrontal and orbito-frontal cortices during the last part of the counting (i.e., over a 2000 ms interval before the platform translation) were greater in the dual task than in the single task (Fig. 6B). This confirmed the participants’ engagement in the cognitive task (McGuire & Botvinick, 2010; Wallis, 2007). Note that no such activities of the frontal lobe were observed in the 2 participants that were discarded from the analyses, due to their high error rate.

##### Postural reaction in response to surface translation

The ANOVA indicated that postural reaction was of greater magnitude when performing the dual task (combined cognitive and motor tasks) than the single motor task (F_1,18_= 4.78; p=0.04), and when standing on the biomimetic as compared to the grooved surface (F_1,18_= 5.92, p=0.03) (Fig. 6C). No significant interaction was observed between Task and Surface (F_1,18_= 0.76; p=0.39). The ANOVA also revealed that the duration of the postural reaction was shorter when performing a dual task (F_1,18_= 21.183; p < 0.05) and when standing on the biomimetic surface (F_1,18_= 6.95; p = 0.02) (Fig. 6D). The interaction Task x Surface was not significant (F_1,18_= 0.43; p=0.51).

Although the ANOVAs performed on the variables related to the postural reaction did not reveal significant Task x Surface interactions, the inspection of the results shown in Fig. 6 suggests that the most efficient postural reaction was found when participants stood on the biomimetic surface in the dual-task condition (i.e., greater amplitude (Fig. 6C) and smaller duration (Fig. 6D) of the postural reaction). In support with this assumption, the results of planned contrasts post-hoc tests revealed that the postural reaction had significantly greater amplitude (F = 10.51; p < 0.005) and smaller duration (F = 37.70; p < 0.001) when standing on a biomimetic surface, rather than in the 3 other conditions.

## Discussion

The intriguing result of *Experiment 2* is that the postural reaction observed during the platform translation had shorter duration and greater magnitude in the dual task than in the single task, irrespectively of the standing surfaces. These behavioral features comply with the spatiotemporal characteristics of an efficient postural reaction (Ikai et al., 2003; Horak et al., 1997 for a review). They were observed to a greater extent when the participants stood on the biomimetic surface. Therefore, the participants’ engagement in the cognitive task did not have a deleterious consequence on the postural control as often reported in previous studies (e.g., Shumway-Cook et al., 1997; Kerr et al., 1985; Rankin et al., 2000; Melzer et al., 2001). The greatest efficiency of the postural reaction observed in participants standing on the biomimetic surface could stem from a greater capacity to shift the attentional focus from the primary motor task to the secondary cognitive task. Such external focus is known to diminish motor-related conscious attentional processes (Vuillerme & Nafati, 2007; Wulf & Prinz, 2001, for a review) compared to an internal focus of attention (Sherwood et al., 2020), and improve motor performance. This has been clearly demonstrated by Wulf et al. (1998) in a study in which participants had to make oscillatory movements (ski-type slalom movements) when standing of platform mounted on wheels that ran laterally on two bowed rails. Elastic rubber belts attached to the platform ensured that the platform returned to the center position during the oscillatory movements. The authors showed that the motor performance decreased when the participants’ attention was focused on the force that the feet should exert on the supporting platform (i.e., internal focus) as compared to when their attention was focused on the wheels of the platform (external focus). These results, combined with those of *Experiment 1* showing similar postural reactions between the different surfaces, suggest that during small accelerations of the standing platform, the advantage of standing on a biomimetic surface to safeguard stability is expressed when one is involved in a dual task (*Experiment 2*). Although it is often the case that one is engaged in a cognitive task while standing (e.g., listening to people, singing while showering, etc.), greater platform accelerations could be needed for the biomimetic surface to improve postural reactions compared to other types of surface.

It is possible that the equilibrium demands in response to support translations decreased when standing on a biomimetic surface, as also suggested by the smaller theta power observed in *Experiment 1* with this surface. The biomimetic surface may therefore facilitate the use of low-level sensorimotor loops, which are less permeable to cognitive load and which enable speedy performance (see Wulf & Prinz, 2001 for a review). As mentioned in Discussion 1, thalamic projections to the left pre-cuneus (Cunningham et al., 2017; Gamberini et al., 2020), which has dense interconnections with the motor and premotor cortices (see Krubitzer et Disbrow 2008 for a review) may have contributed to the facilitation of the neural responses to the tactile stimulation observed with the biomimetic surface (i.e., increased P1N1 SEP). These thalamocortical connections areas could constitute the neural underpinning of the efficient spatiotemporal pattern of the postural reaction when standing on the biomimetic surface.

## Acknowledgement

We thank Frédéric Pous (Institut des Sciences du Mouvement, Marseille) for his assistance in the fabrication of the supporting surfaces

